# Differential Effectiveness of Poloxamer 188 in Dystrophic Cardiomyopathy Reveals Complex Pathophysiology

**DOI:** 10.1101/2024.09.04.611221

**Authors:** Jackie Stevens, F. Parker Banyard, Danielle Millner Balagtas, Derek Williams, DeWayne Townsend

## Abstract

Synthetic membrane stabilizers, such as poloxamer 188 (P188), have great promise in the treatment of Duchenne muscular dystrophy and other diseases associated with disrupted membranes. The aim of this study was to assess the efficacy of P188 to limit myocardial damage in models of muscular dystrophy subjected to highly injurious stress. These studies use male and female mdx and β-sarcoglycan null (β-SG^-/-^) mice subjected to an isoproterenol stress protocol in the presence or absence of P188. We show that P188 provides a significant level of protection to male mdx hearts from the isoproterenol induced injury. Surprisingly, we find that P188 has no protective actions in female mdx mice or in β-SG^-/-^ mice of either sex. These results suggest that the mechanism of action of P188 is more complicated than previously thought. The presence of sex differences in the absence of dystrophin is not consistent with a model of P188 working by biophysical restoration of membrane integrity. Furthermore, the absence of protective effects of P188 in β-SG^-/-^ myocardium indicates that there are important differences in the mechanisms of membrane damage in hearts lacking dystrophin and hearts lacking the sarcoglycan complex. Developing a better understanding of these differences will be important in the development of therapies for dystrophic cardiomyopathy and other forms of heart disease.

## Introduction

The muscular dystrophies represent a group of genetic diseases that affect striated muscle and are characterized by muscle weakness and the loss of contractile tissue. Many of the most serious muscular dystrophies are associated with a complex of proteins associated with the membrane of striated muscle cells. The first member of this complex to be identified was dystrophin, a large cytoskeletal protein whose absence results in the X-linked Duchenne muscular dystrophy (DMD) (1). Shortly thereafter many additional proteins were found to associate with dystrophin (2–5), many of which were subsequently identified as loci for other muscular dystrophies (6–10). The sarcoglycan complex consists of four membrane bound glycoproteins that are tightly associated with dystrophin. The absence of any member of the complex causes significant reductions in all other components of the complex and results in severe muscle diseases, collectively called sarcoglycanopathies.

In both DMD and the sarcoglycanopathies in addition to skeletal muscle wasting there are significant deficits in cardiac function (11–17). The clinical significance of dystrophic cardiomyopathies are becoming more apparent with the improvements in the management of the respiratory complications associated with these diseases. There are promising potential therapies on the horizon, but there are significant concerns about the efficacy of these new therapeutic approaches in the heart. Genetic mouse models have been invaluable in developing our understanding of the pathophysiology of dystrophic cardiomyopathies and have been essential in the development of therapeutic approaches now entering the clinics. However, the disease that these mice present is relatively mild compared to the severe disease evident in patients with muscular dystrophy. One successful approach for the evaluation of a therapy’s efficacy is to measure its ability to prevent disease in animal models that have been stressed; in studies of skeletal muscle this often takes the form of mechanical stresses such as lengthening contractions. In the heart there are no universally accepted stressors with approaches ranging from genetic stress, in which other genes are partially or completely removed, aging stress, or workload stress (18–20). This latter approach has several advantages in that it can be performed quickly, and like lengthening contractions, it provides a significant challenge to dystrophic muscle allowing therapies with partial efficacy to be identified.

The similar clinical presentation and muscle pathology coupled with their biochemical association suggest that DMD and the sarcoglycanopathies may share a common pathophysiology. However, several studies suggest that there may be some important differences (21). In this study we examine the efficacy of poloxamer 188 (P188) to provide protection against large increases in cardiac workload induced by isoproterenol. Previous studies have demonstrated that P188 protects the heart of both the mdx mouse and the golden retriever model of DMD (22,23). However, these studies examined the dystrophic heart following minutes of increased workload or longer periods of baseline contractile function.

Here we use a single isoproterenol bolus injection in two mouse models of muscular dystrophy to generate several hours of increased contractile activity which represents a significant stress in the dystrophic heart and resulting in considerable myocardial injury.

## Methods

### Animals

The mice used in these studies were obtained from colonies maintained at the University of Minnesota. The mdx mice were derived from mice purchased from Jackson Laboratories line C57BL/10ScSn-DMDmdx, new breeding stock were introduced into the colony every 4-5 generations. The β-sarcoglycan (β-SG^-/-^) mice were obtained from a colony derived from mice obtained from Jackson Laboratories line B6.129-Sgcb^tm1Kcam^/1J and bred to homozygosity. All animals were 4-6 months of age. All experimental protocols were approved by the University of Minnesota Institutional Animal Care and Use Committee.

### Drug Treatments

P188 was dissolved in 0.9% NaCl at a concentration of 150 mg/ml and filter sterilized. Mice were injected subcutaneously with 460 mg/kg P188 at least 1 hour prior to injection of isoproterenol. (-)-Isoproterenol hydrochloride (Iso; Sigma #I6504) was dissolved in 0.9% NaCl at a concentration of 6 mg/ml. This solution was filter sterilized and stored protected from light at 4°C. Single 10 mg/kg intraperitoneal injection of isoproterenol was performed between 8:00 and 10:00 AM. Hearts were harvested 30 hours later, arrested in 60 mM KCl in PBS, embedded in OCT, and flash frozen in liquid nitrogen cooled isopentane. Blocks were stored at -80 °C until sectioned. Hearts were sectioned with 7 µm thick sections being used for analysis.

### Injury Assessment

Sections were blocked with 10% goat serum and stained with wheat germ agglutinin (WGA) conjugated to Alexa-647 (ThermoFisher #W32466, 5 µg/ml) and goat anti-mouse IgG-594 (H + L) secondary antibody (ThermoFisher #R37121) for 1 hour at room temperature. Sections were washed three times with PBS for 5 minutes each prior to the application of ProLong Gold Antifade Mounting media with DAPI (ThermoFisher #P36941).

Images were collected using a Keyence BZ-X800E microscope with the 10x objective and automated imaging of DAPI, endogenous IgG, and WGA fluorescent signals. Montages of the individual images were generated and used for subsequent analysis. Intracellular IgG staining was measured by thresholding an image resulting from the subtraction of the WGA channel from the IgG channel. This procedure effectively removed any extracellular IgG staining allowing for the efficient identification of intracellular IgG, representing regions in which membrane integrity had been disrupted. Images were thresholded and pixels quantified using ImageJ. Similarly, the pixel counts for the entire myocardium were determined. Total myocardial injury was reported as the percentage of myocardium that was positive for intracellular IgG. Analyses were performed by investigators blinded to sex or treatment of the images being analyzed. Note that the WGA signal was collected in the infrared spectrum, but is displayed as green in the figures to maximize contrast.

### Statistical Analysis

Injury data were compared by unpaired T-tests examining the effect of P188 pretreatment using the statistical analysis software R(24).

### Data Availability

The data underlying the results presented here are available from the corresponding author upon request.

## Results

Previous work demonstrated that P188 administered IV provided protection to the dystrophic heart at rest and during stress (22,23). Additional work demonstrated that subcutaneous (SQ) administration of P188 provided protection to the skeletal muscle of mdx mice (25). This work is consistent with the rapid prolonged systemic distribution of P188 following SQ administration (26). To evaluate the capability for SQ delivered P188 to protect the mdx heart under periods of stress, we administered 460 mg/kg P188 via a SQ injection 1 hour prior to intraperitoneal (IP) injection of 10 mg/kg isoproterenol. This dose of isoproterenol has been demonstrated to cause significant levels of cardiac injury in male mdx mice (20). Male mdx mice administered a SQ injection of saline displayed an extensive myocardial injury as measured using IgG incorporation (Fig. 1). In contrast, male mdx mice receiving P188 showed a significant reduction in the level of myocardial injury, as measured by intracellular IgG. To assess the presence of any sex differences in the susceptibility of mdx mice to injury, we performed a similar experiment with female mdx mice. Thirty hours following isoproterenol injection, female mdx mice have levels of myocardial injury similar to that observed in male mdx mice (Fig. 2). Surprisingly, female mdx mice pretreated with P188 had the same level of workload-induced myocardial injury as saline treated mice.

**Figure 1.**
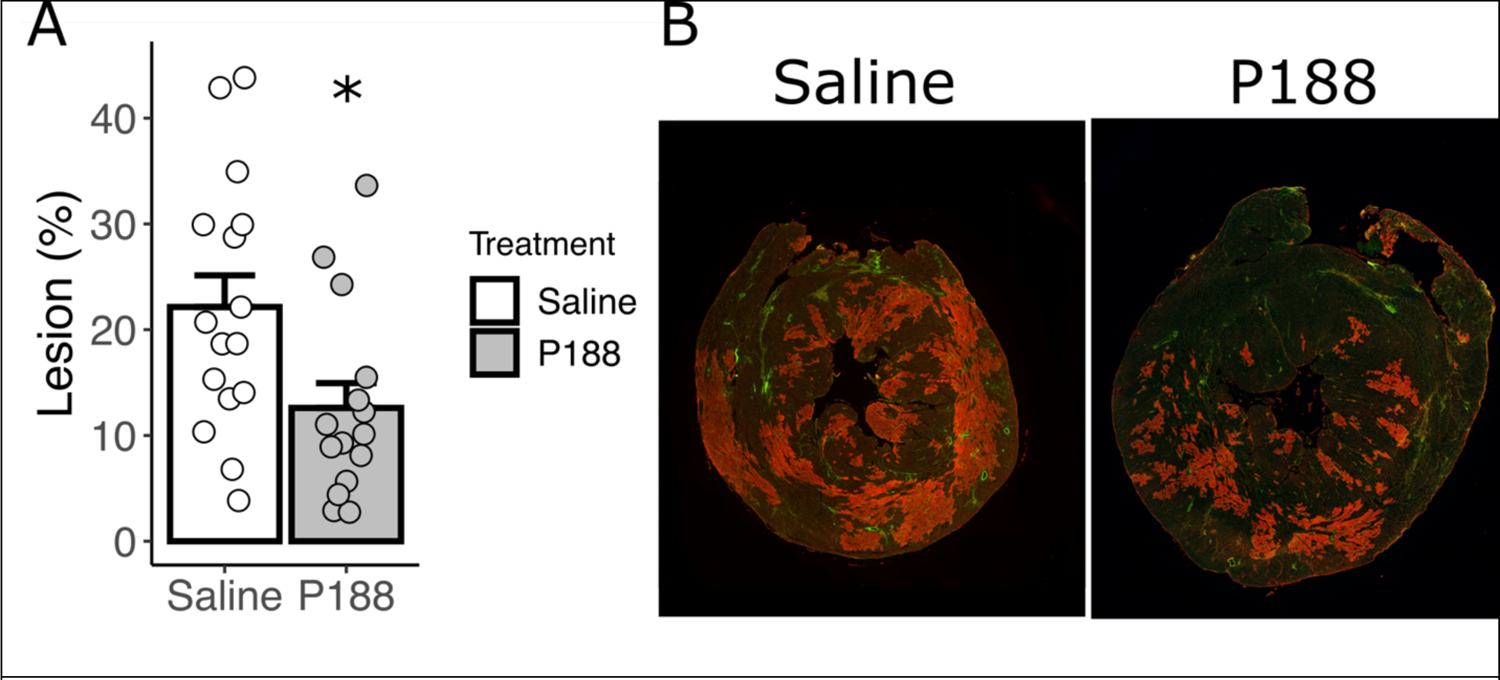
P188 pretreatment significantly reduced workload-induced myocardial injury in male mdx mice. Pretreatment of male mdx mice with 460 mg/kg via SQ injection 1 hour prior to injection with 10 mg/kg isoproterenol significantly reduced the workload-induced injury detected 30 hours after the isoproterenol injection. * - p = 0.018; n=15-16.

**Figure 2.**
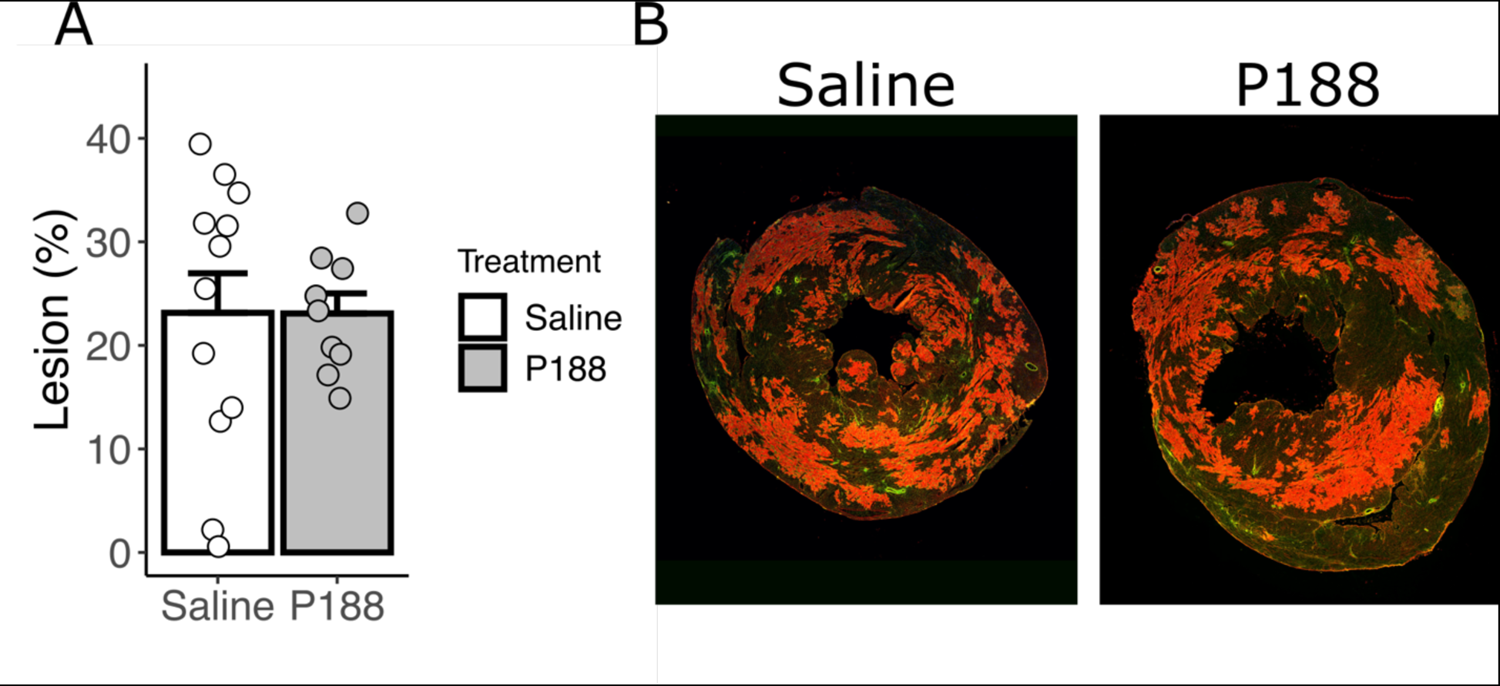
P188 pre-treatment has no significant effect on myocardial injury 30 hours after isoproterenol injection in female mdx mice. In contrast to male mdx mice, the hearts of female mdx mice were not protected from workload-induced injury by the presence of P188. n= 9-12

Patients with sarcoglycanopathies, especially those with mutations in β-, γ-, or δ-sarcoglycan genes, develop significant cardiomyopathy that is clinically similar to that observed in DMD patients (13–17). Both are characterized by progressive heart disease with declines in contractile function and prominent myocardial fibrosis present in advanced stages. Mouse models lacking β-, γ-, or δ-sarcoglycan display evidence of membrane disruption similar to that observed in the mdx mouse (27–31). The presence of breaks in membrane integrity suggested that treatment with the membrane sealant P188 would be of potential therapeutic benefit.

Mirroring the studies performed in mdx mice, P188 was provided in a SQ injection to male mice lacking β-sarcoglycan. Following this pretreatment, male β-sarcoglycan null (β-SG^-/-^) mice were injected with 10 mg/kg isoproterenol. Much like male mdx mice, the hearts of male β-SG^-/-^ mice displayed widespread injury throughout the myocardium (Fig. 3). In contrast to mdx mice, the workload-induced injury in the male β-SG^-/-^heart was not reduced by pretreatment with P188. Female β-SG^-/-^hearts are relatively resistant to isoproterenol-induced myocardial injury (32) and the injury present in these hearts was also not altered by P188 pretreatment (Fig. 4).

**Figure 3.**
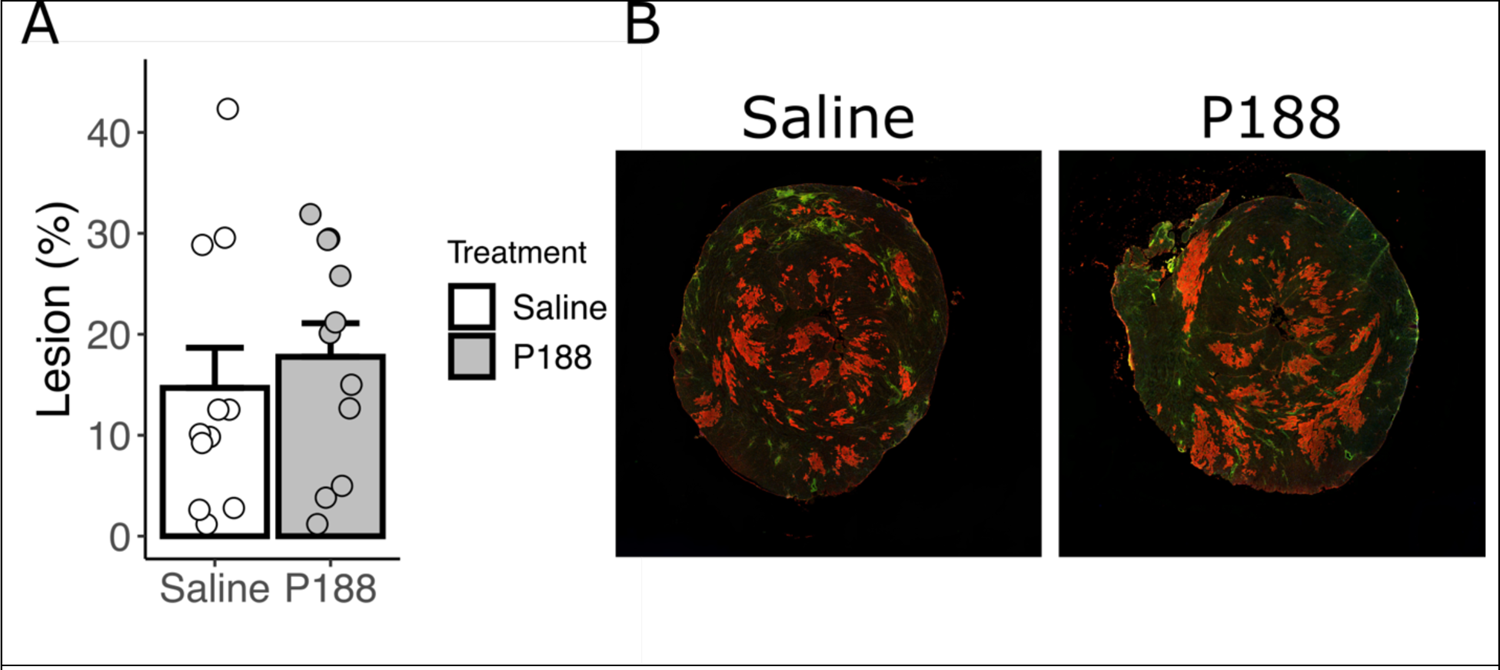
Male β-SG^-/-^ mice are susceptible to workload-induced injury, but this injury is not prevented by P188 pretreatment. Measurement of myocardial injury 30 hours following injection of 10 mg/kg isoproterenol. n= 11

**Figure 4.**
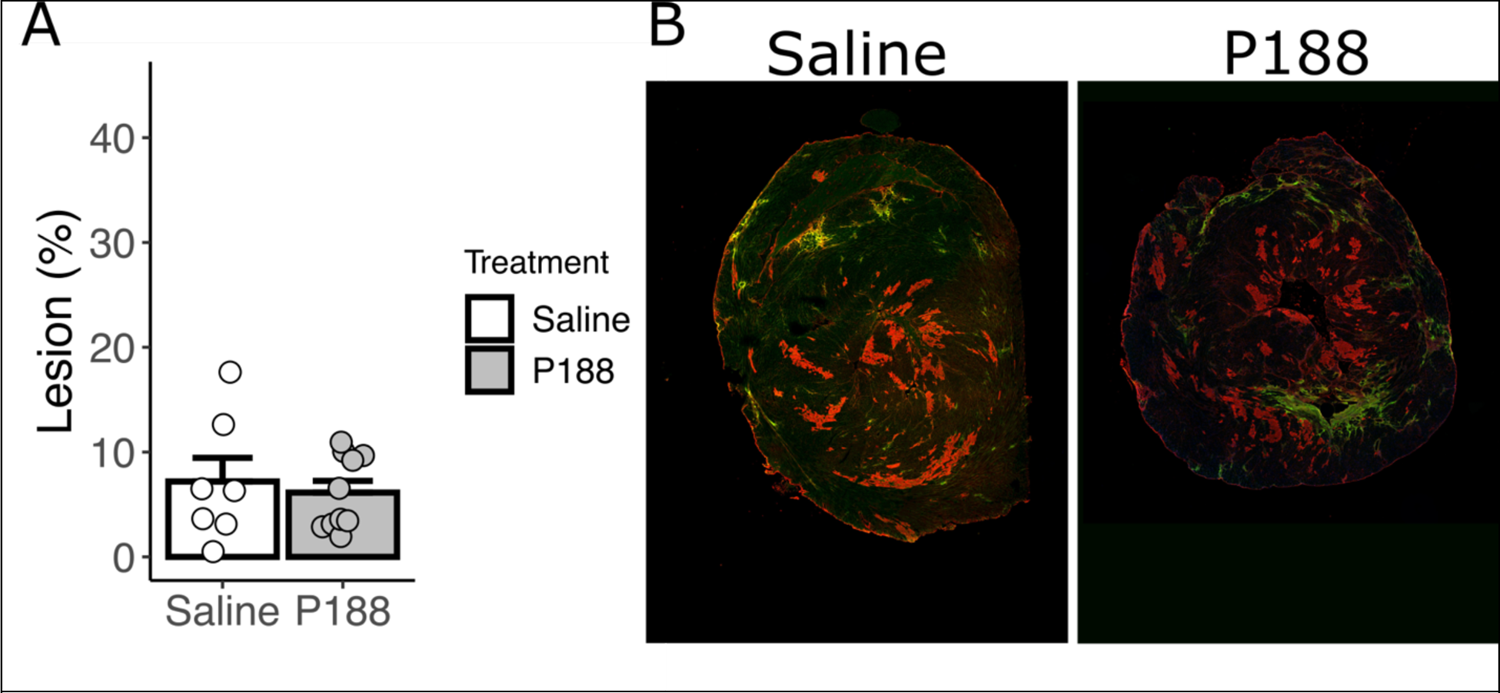
Female β-SG^-/-^ mice are relatively resistant to workload-induced injury and what injury they have is not affected by P188 pretreatment. Measurement of myocardial injury 30 hours following injection of 10 mg/kg isoproterenol. n= 7-10

## Discussion

Poloxamer 188 (P188) is a synthetic membrane stabilizing agent that has shown promise as a therapy in animal models of DMD, by providing protection to both the heart and skeletal muscle (22,23,25,33). The details of the mechanism of this protection are currently poorly understood, but there is a consensus that P188 protective actions derive from its interaction with the plasma membrane of striated muscle (34–38). Biophysical and in silico studies have demonstrated that P188 may function to stabilize membranes under lateral traction (36,39). The sex-difference observed here suggest that the pathophysiology of dystrophin loss is different between male and female mdx mice and that P188 is only able to disrupt the pathophysiology in male mdx mice. Prior studies in the mdx mouse have either shown no sex-difference (21,40) or indicate that female mdx mice have worse cardiac function (41). The results of this study also appear to contradict the studies of Spurney et al. which observed a protective action of P188 in female mdx subjected to isoproterenol challenge (33). In this prior study, the severity of the injury produced was very modest likely related to the lower doses of isoproterenol required for survival of mdx mice following osmotic pump implantation. Our data provides evidence that with highly injurious doses of isoproterenol, female mdx mice are not protected by doses of P188 that do provide protection to male mdx mice.

The inability of P188 to alter the degree of myocardial injury in β-SG^-/-^ hearts suggests that there are significant differences in the pathogenic mechanisms between these two models of muscular dystrophy. Several previous studies provide evidence supporting this conclusion. Our prior work demonstrated differences in passive tension of isolated myocytes and myocardial fibrosis between mdx and γ-SG^-/-^ and δ-SG^-/-^ hearts (21). The presence of a unique pathogenic mechanism for the sarcoglycanopathies is further supported by the worsening phenotype of mdx mice lacking δ-SG (42). Finally, the distinct sex differences between β-SG^-/-^ and mdx models of dystrophic cardiomyopathy another pathological difference between these two mutations (32); although the impact of genetic background may also contribute to these differences.

The absence of a therapeutic effect of a pharmaceutical agent can result from a variety of causes. For example, differences in metabolism or distribution may alter the final concentration present within the tissue of interest, thus affecting its efficacy. This principle has been documented with P188 where the efficacy of which is highly dependent on the nature of the administration with SQ administration providing protection to skeletal muscle, but IP and IV administration were found to be ineffective (25). This dynamic appears to result from the rapid clearance of P188 from the plasma following IV administration, but it is slowly absorbed from the SQ injection site (26). It is notable that studies in rats and dogs did not identify any sex-differences in the metabolism or pharmacokinetics of P188 following IV administration (43,44). Together this data argues against differences in the metabolism or distribution of P188 being responsible for the differential efficacy observed in these studies.

In summary, we observe that P188 is selectively protective of male mdx mice. Indicating that the absence of dystrophin has unique pathological characteristics in males that are particularly amenable to P188 treatment. These results argue against a model of P188 functioning as a simple membrane sealant. The mechanism by which P188 provides therapeutic benefit to male, but not female, mdx mice is unclear. P188 has been shown to augment membrane repair processes (34), which may be differentially important across sexes; alternatively, P188 has shown efficacy in ischemia reperfusion injury (45,46), perhaps reactive oxygen species generation is more pronounced in male mdx hearts. Overall, these studies highlight the complexities of the pathophysiology of the muscular dystrophies and their potential therapies.

